# Beat-Relevant Signals in Auditory Cortical Responses to Musical Excerpts

**DOI:** 10.1101/481473

**Authors:** Vani G. Rajendran, Nicol S. Harper, Jan W. H. Schnupp

## Abstract

Musical beat perception is widely regarded as a high-level ability involving widespread coordination across brain areas, but how low-level auditory processing must necessarily shape these dynamics, and therefore perception, remains unexplored. Previous cross-species work suggested that beat perception in simple rhythmic noise bursts is shaped by neural transients in the ascending sensory pathway. Here, we found that low-level processes even substantially explain the emergence of beat in real music. Firing rates in the rat auditory cortex in response to twenty musical excerpts were on average higher on the beat than off the beat tapped by human listeners. This “neural emphasis” distinguished the perceived beat from alternative interpretations, was predictive of the degree of consensus across listeners, and was accounted for by a spectrotemporal receptive field model. These findings indicate that low-level auditory processing may have a stronger influence on the location and clarity of the beat in music than previously thought.

## Introduction

The perception of a steady pulse or beat in music is a curious phenomenon that arises from the interaction between rhythmic sounds and the way our brain processes them. There are two things that make musical beat perception particularly intriguing. Firstly, no mammalian species apart from humans consistently show spontaneous motor entrainment to the beat in music (e.g. tapping a foot, nodding the head, moving the body)^1–4^. Secondly, despite beat being a subjective percept rather than an acoustic feature of music, individual listeners tend to overwhelmingly agree on where the beat is. Some of this consistency might be due to certain “top-down” constraints such as cultural and cognitive priors^5–7^. However, apart from theory^8,9^, relatively little is known about the neurophysiological dynamics that cause the feeling of musical beat to emerge in the first place.

A key piece of information currently lacking is which aspects of the neural representation of music might be important for the induction of beat. Previous cross-species work revealed that firing rates as early as the auditory midbrain are significantly higher on the beat than off the beat in simple rhythms constructed from identical broadband noise bursts^10^. If large firing rate transients resulting from low-level auditory processing are indeed necessary for the induction of beat, then this insight could shed light on the dynamics of the entrainment of cortical oscillations to beat^11–16^, the role played by the motor system^8,17–26^, and why different species differ so much in their beat perception and synchronization capacity^27^.

Importantly, if a consequence of auditory processing is to create points of neural emphasis that predispose beats being felt there, then we should observe this not just for simple rhythmic “laboratory sounds,” but also for real music. Twenty musical excerpts^28^, which were diverse in tempo and musical genre, were played to three anesthetized rats while recording extracellularly from auditory cortex. In line with previous findings, population firing rates were higher on the beat than off the beat, and large on-beat to off-beat firing rate ratios were a distinguishing feature of the consensus beat interpretation across human listeners. Comparison with the output of an auditory nerve model revealed that small effects may already be present at the auditory periphery but are amplified substantially in cortical responses. Musical excerpts that evoked a larger cortical on-beat emphasis also showed a stronger consensus in tapping behavior across listeners. Finally, these results could be accounted for by the spectrotemporal receptive field properties of recorded units. These findings add to growing evidence that beat perception is not entirely culturally determined, but is also heavily constrained by low-level auditory processing common to mammals.

## Results

Neural activity from a total of 98 single and multi-units were analyzed in response to 12 repeats of the first 10 seconds of 20 musical excerpts taken from the MIREX 2006 dataset online, which included beat annotations made by 40 human listeners^28^. In all songs, listeners reported a steady beat well within the first 10 s. The most common tapping pattern for each excerpt was taken to be that excerpt’s “consensus” beat interpretation (see *Methods*), and consensus tapping rates ranged from 0.7 Hz to 3.7 Hz (42 to 222 beats per minute, corresponding to beat periods of 1.42 down to 0.27 s). The analyses that follow investigate correspondences between firing rates in the rat auditory cortex around the consensus beat as reported by human listeners.

### Auditory cortical firing rates are higher on the beat than off the beat

For each song, the 100 ms time window following each consensus tap was defined as on-beat, and all time excluding these on-beat windows was defined as off-beat (the results are not sensitively dependent on this precise definition, see *Methods*). Fig 1A shows the average on-beat population firing rate plotted against the off-beat population firing rate for each of the 20 tested musical excerpts. On-beat firing rates were significantly larger than off-beat firing rates (p<10^−4^, Wilcoxon paired signed-rank test, N=20 songs), an observation that is consistent with previous work examining gerbil midbrain responses to simple rhythmic patterns^10^. The beat-triggered average population firing rate in the 200 ms window around consensus beats (averaged across all beats in all excerpts) provides a more detailed picture of population neural activity around the beat (Fig 1B). The distribution of on:off-beat ratios (OORs; average on-beat firing rate divided by average off-beat firing rate) for each recorded unit (N=98) is shown in Fig 1C. An OOR > 1 indicates that firing rates were higher on the beat than off the beat. Most units show an OOR > 1, and the bimodal distribution suggests that there may exist distinct sub-populations in the recorded data, one with OORs centered around 1 and the other with OORs around 1.5.

**Fig 1.**
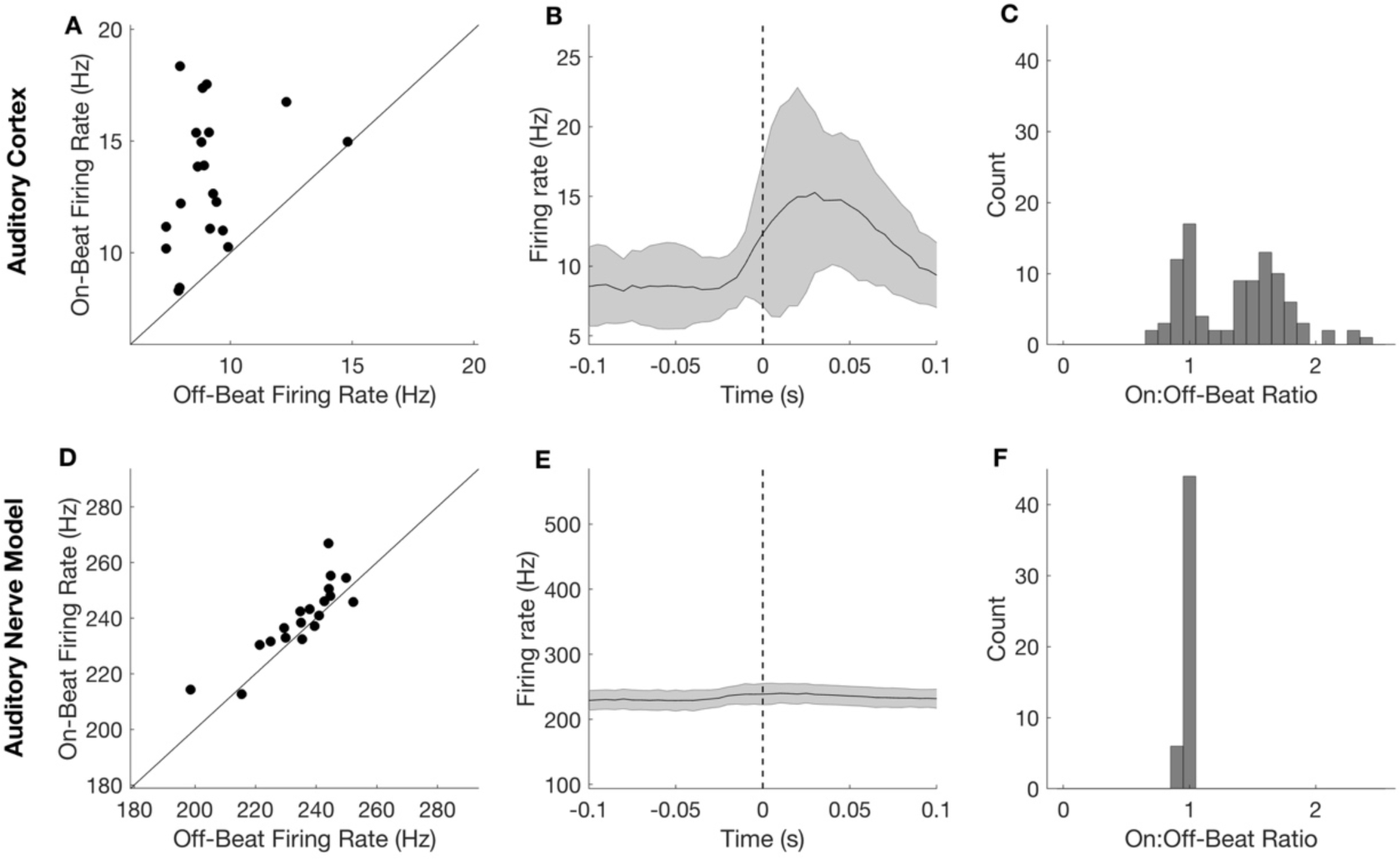
Consensus beat-triggered neural activity and on:off-beat ratios (OORs) in the auditory cortex and auditory nerve. **(A)** Mean on-beat versus off-beat population firing rate in auditory cortical neurons. Each dot is one musical excerpt. On-beat firing rates are significantly higher than off-beat firing rates (p<10^−4^, Wilcoxon paired signed-rank test, N=20 songs) **(B)** Population “beat-triggered” average firing rate in the auditory cortex in a 200 ms window around the consensus beat times ± standard deviation across the 20 musical excerpts. **(C)** Histogram of on:off-beat firing rate ratios (OORs) for each recorded unit (N = 98), where “on-beat” is the average firing rate during the 100 ms post-tap window, and “off-beat” is the average firing rate over the entire song excluding on-beat windows. **(D)** Same as A, but for population activity based on an auditory nerve model with 50 log-spaced frequency channels between 150 Hz and 24 kHz. Predicted firing rates at the auditory nerve were significantly higher on the beat than off the beat (p<0.005, Wilcoxon paired signed-rank test, N=20 songs) **(E)** Same as B, but for population activity based on the auditory nerve model. **(F)** Same as C, but for auditory nerve model fibers (N=50).

For comparison, an auditory nerve model^29^ was used to predict firing rates at the auditory nerve for 50 logarithmically spaced frequency channels between 150 Hz and 24 kHz. Fig 1D shows predictions of on-beat versus off-beat population activity at the auditory nerve. Notably, the auditory nerve model would also predict higher average population firing rates on the beat than off the beat (p<0.005, Wilcoxon paired signed-rank test, N=20 songs). Fig 1E–1F show beat-triggered averages and OORs for auditory nerve model fibers. OORs based on the auditory nerve model, though significantly larger than one, are much smaller than cortical OORs (p<10^−4^, Wilcoxon paired signed-rank test, N = 20 songs).

### A large neural emphasis is a distinguishing feature of the consensus beat

While we have shown that firing rates are higher on the beat than off the beat, this on its own does not imply that large OORs are necessarily relevant to beat perception. From a purely signal processing perspective, a musical excerpt could theoretically be perceived as having any combination of tempo and time signature, and if most of these possible alternative beat interpretations were associated with more or less equally large OORs, then large OORs would be of little value as physiological markers of musical beat. Therefore, if a large OOR is relevant for the induction of beat, we hypothesized that it should be large for the consensus beat relative to plausible alternatives.

To test this, we computed hypothetical OORs for the full range of plausible beat period and phase combinations. For each song, possible beat periods (representing the different rates at which a listener might tap) were allowed to range from 0.2 s to 2 s (5 Hz down to 0.5 Hz) sampled in 20 ms steps. Likewise, for each beat period, the phase offset was allowed to range from 0 up to the full beat period sampled in 20 ms steps to capture the fact that two listeners tapping at the same rate may nonetheless exhibit different interpretations of the beat if their taps, rather than being synchronous, have a constant offset between them. The OOR was then computed for each of these beat interval and beat offset combinations, resulting in 4,995 possible OOR values for each musical excerpt. The heatmaps in Fig 2A and 2B show the computed set of plausible OOR values calculated from cortical and auditory nerve model firing rates, respectively, for an example musical excerpt, with possible beat periods on the y-axis and possible starting phase offsets on the x-axis (see Supplementary Figs S1-S2 for heatmaps of all musical excerpts). The histograms in Fig 2C and 2D pool together hypothetical (in gray) and consensus (in red) OOR values from all musical excerpts (histograms for individual excerpts in Supplementary Fig S3-S4).

**Fig 2.**
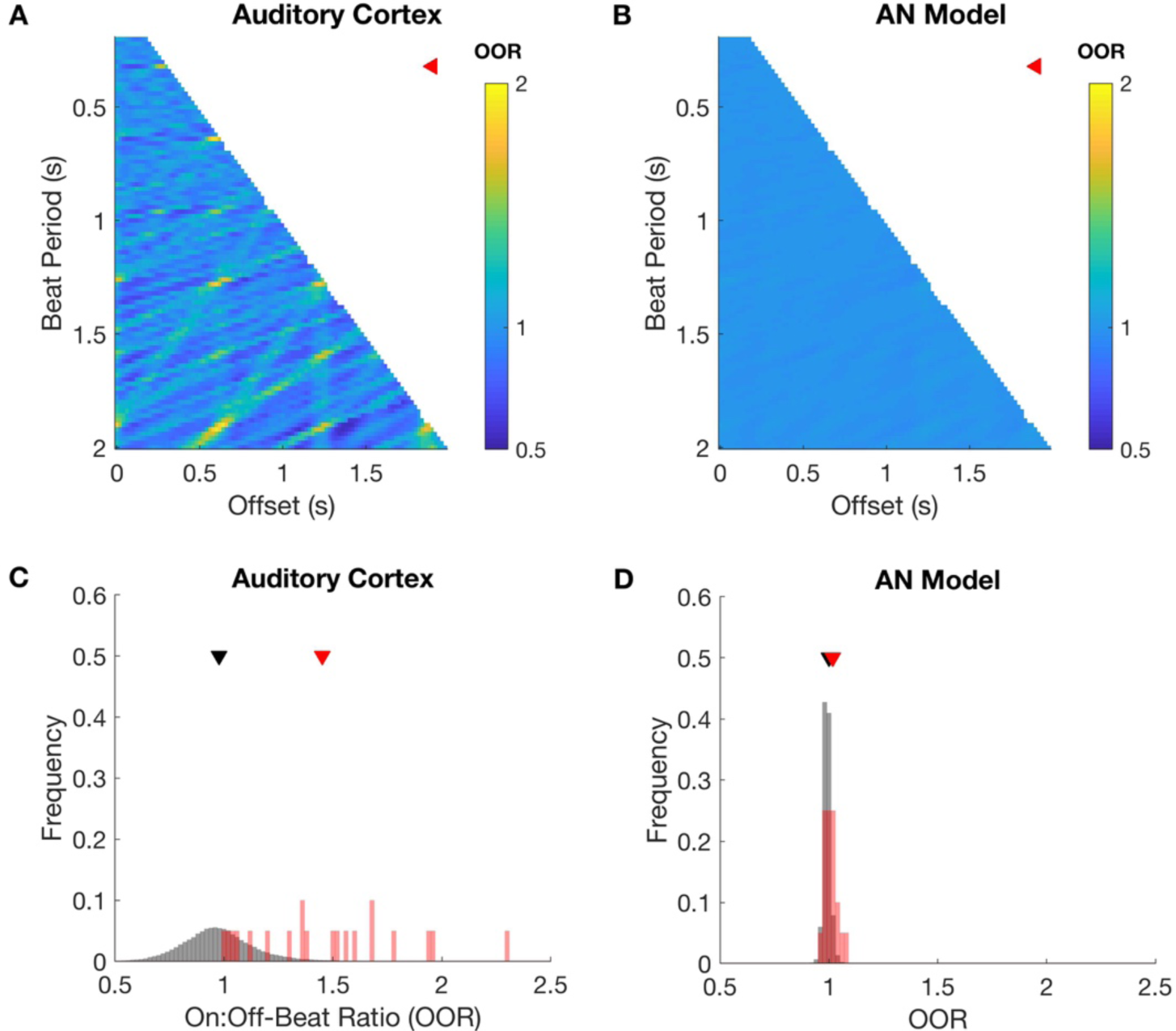
How does the consensus beat compare with other possible beat structures? **(A)** Heatmap depicting cortical on:off-beat ratios for plausible beat period (y-axis) and beat phase offset (x-axis) combinations between 200 ms and 2 s (or tap rates of 5 Hz down to 0.5 Hz) for one example musical excerpt. Color indicates the OOR value. **(B)** Same as A, but for population activity based on the auditory nerve model. **(C)** Histogram pooled across musical excerpts of all OOR values (gray), and consensus OOR values (red) in the auditory cortex. **(D)** Same as C, but based on OOR values from the auditory nerve model.

If the OOR is a distinguishing feature of the perceived beat, we would expect it to rank above the 50^th^ percentile of the underlying distribution of hypothetically plausible OORs for a given musical excerpt. As hypothesized, the consensus OORs rank significantly larger than the 50^th^ percentile, both in the auditory cortex (p<10^−4^, Wilcoxon signed-rank test, N = 20 songs), and in the auditory nerve model (p<0.005). However, the percentiles were significantly larger in the auditory cortex than in the auditory nerve model (p<0.005, Wilcoxon paired signed-rank test, N = 20 songs). Notably, 14 out of the 20 musical excerpts tested had consensus OORs above the 95^th^ percentile in the auditory cortex, in contrast to only 7 out of 20 based on the auditory nerve model. Additionally, fewer hypothetical beat interpretations resulted in large OORs in the auditory cortex, as evidenced by the higher skewness, or longer right tails, of the OOR distributions in the auditory cortex compared to those based on the auditory nerve model (p<10^−4^, Wilcoxon paired signed-rank test, N = 20 songs). Together, these results suggest that a large OOR is a feature that distinguishes the consensus beat from most other possible beat structures, and that two important consequences of auditory processing might be an amplification of small differences in OOR already present at the auditory periphery, and a further restriction of the candidate beat interpretations that would result in large OORs.

### The stronger the on-beat neural emphasis, the stronger the tapping consensus

It is clear from Fig 2C (and Supplementary Fig S3) that consensus OORs are consistently among the largest possible OORs across our set of musical excerpts, but they are not always the largest. However, it is not uncommon for the beat in a given piece of music to be perceived in different ways. More often than not, listeners will exhibit a variety of tapping patterns, for example with some tapping twice as fast or half as fast as others, or 180 degrees out of phase with others. Additionally, if the beat is not very salient, there will be uncertainty about when exactly a beat occurs and therefore an increased variance in observed inter-tap-intervals. In such cases, and indeed in the dataset we use, listeners display a range of perceived beat interpretations, and what we have termed the consensus beat is merely the beat interpretation that happens to be favored by a (sometimes narrow) majority of listeners. This variability is illustrated in Fig 3, where for some excerpts tapping behavior was consistent across a large majority of listeners (e.g. Fig 3A), and for others tapping behavior was more variable, indicating a less salient or more ambiguous beat percept (e.g. Fig 3B–3C; see Supplementary Fig S5 and S6 for tapping behavior for all excerpts).

**Fig 3.**
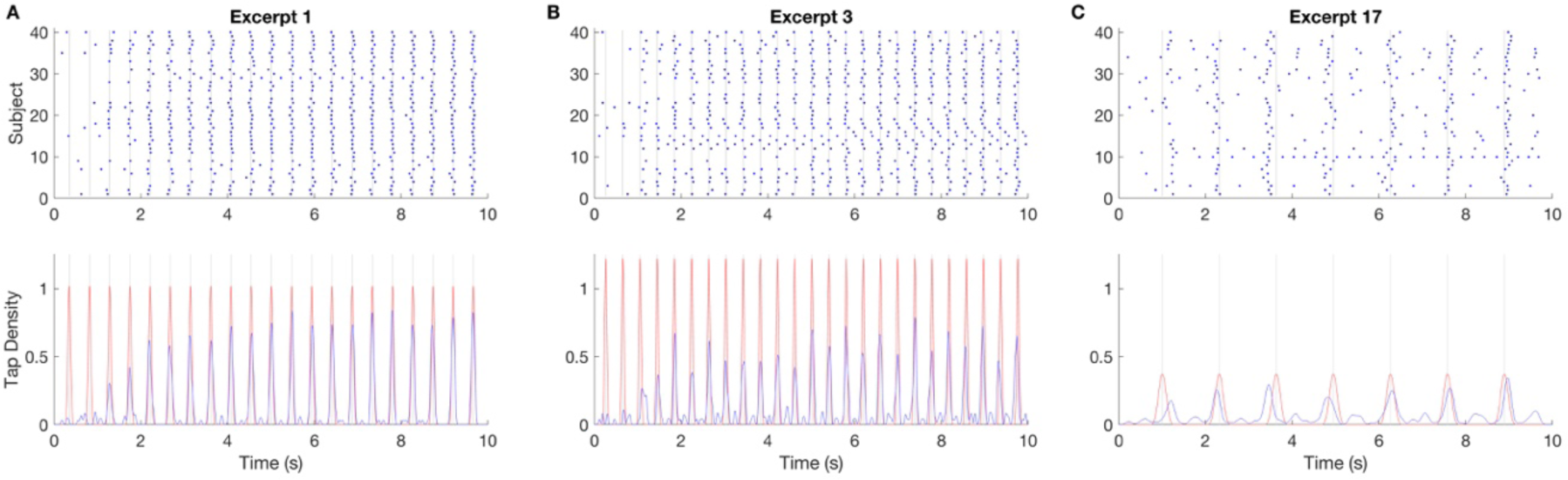
A glimpse into the variability across human listeners tapping to the beat in music. **(A) Top:** Raster plot of tap times for the 40 human annotators across the 10 s excerpt of an example song. Each row is one subject, and location along the x-axis represents when the subject tapped during the 10 s musical excerpt. Consensus beat times are marked by gray vertical lines (see *Methods*). Note that most subjects’ taps line up in time with each other and with consensus beat for this example excerpt. **Bottom:** Tap density estimates based on tap times pooled across subjects, binned with 2 ms bins, and smoothed with a Gaussian kernel with a standard deviation of 5% of the consensus beat period (blue). Shown in red is a smoothed tap density estimate of the “ideal” tap histogram (with realistic motor error) that would have been obtained if all subjects had tapped on every consensus beat (see *Methods*). The correlation between real and idealized density is high for this excerpt (r=0.88), indicating a strong tapping consensus. **(B)** Same as A, but for a musical excerpt with multiple minority beat interpretations and therefore a lower correlation coefficient (r=0.78). **(C)** Same as B, but where the tapping consensus is even weaker (r=0.59). See Supplementary Figs S5 and S6 for all musical excerpts.

However, if we hypothesize that a large OOR predisposes a listener to hear a particular beat interpretation, then we would predict that the excerpts that evoke the largest OORs in cortical responses should also be the ones that evoke the clearest, most unambiguous beat percept across listeners. Can the variability in tapping behavior be explained by the size of OORs in the auditory cortex?

To answer this question, we quantified the strength of the tapping consensus for each song by calculating the correlation coefficient between the smoothed histogram of observed tap times and the smoothed histogram of the “ideal” case in which all 40 listeners would have tapped on each consensus beat within a realistic degree of sensory or motor error (see *Methods*). Examples of observed (blue) and idealized (red) tap density estimates are shown the lower panels of Fig 3.

Consistent with our hypothesis, the size of the consensus OOR evoked in the auditory cortex by a musical excerpt correlated significantly with the strength of the tapping consensus across listeners (Fig 4A; p<0.001, Pearson correlation, N = 20 songs). Neither OOR (p=0.48) nor consensus strength (p=0.44) varied with the consensus tempo of musical excerpts (Pearson correlation, N = 20 songs). Fig 4B and 4C show how OOR and consensus strength, respectively, develop over the course of the 10 s duration of the musical excerpts. Data were split into five 2-s chunks, and OORs and correlation coefficients were calculated based on the data in each chunk. Tapping consensus strength, which is low initially, is nearly at ceiling from about 4 s into the excerpts, indicating that listeners only needed a few seconds to find the beat. OORs, on the other hand, did not change systematically over time, suggesting that the correspondences observed in this study between neural activity and behavior are unlikely to be due to cortical entrainment or buildup in neural responses.

**Fig 4.**
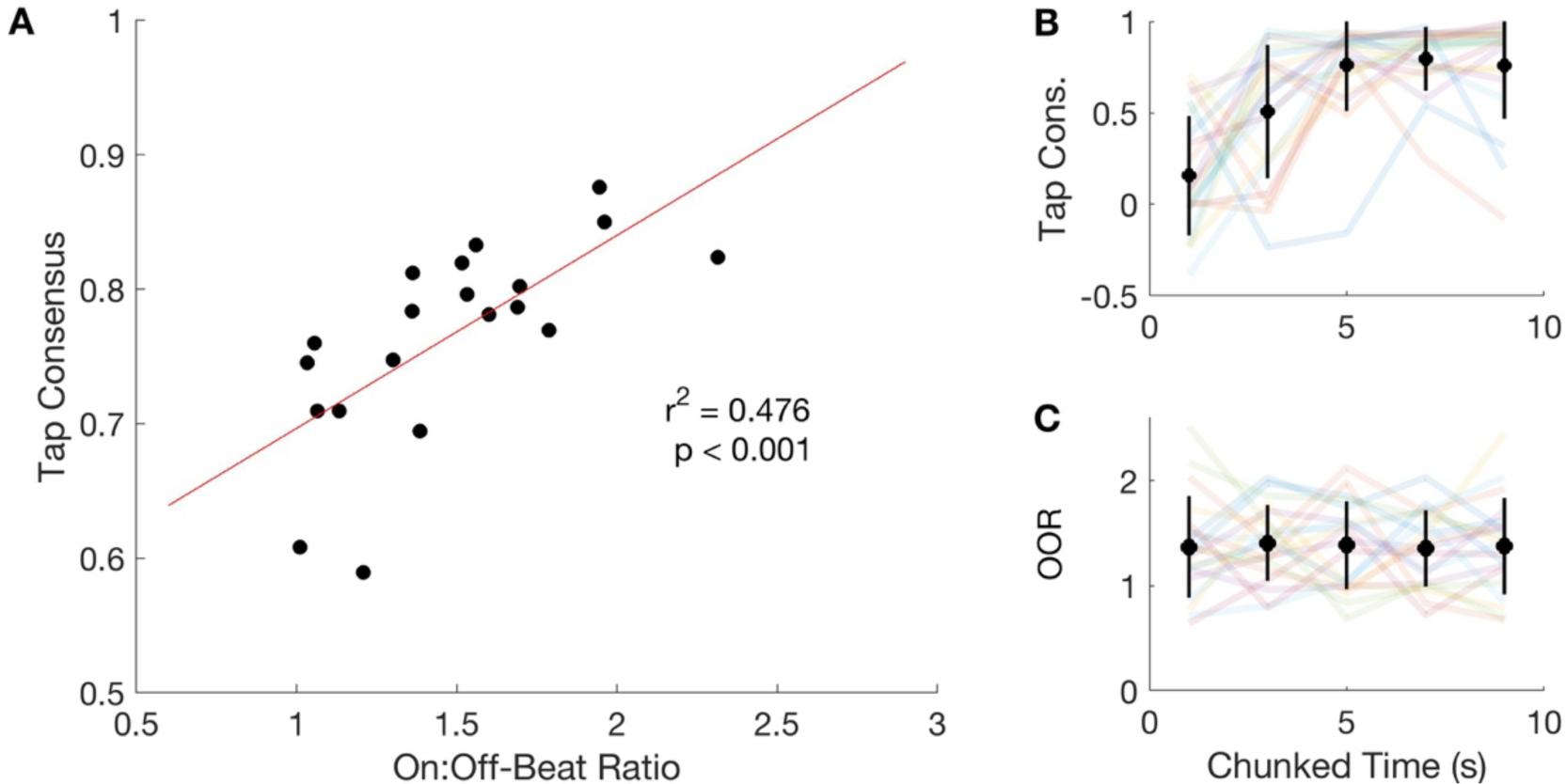
The stronger the on-beat neural emphasis, the stronger the tapping consensus. **(A)** Each dot is one musical excerpt. There is a strong correlation between auditory cortical OOR (x-axis) and the tapping consensus across listeners, quantified as described in Fig 3 (y-axis; p<0.001, Pearson correlation, N = 20 songs). **(B)** Tapping consensus, calculated for each sequential 2 s segment of musical excerpts. Colored lines are individual songs. In black is the mean across songs for each time chunk ± standard deviation. **(C)** Same as panel B but for OOR values.

### Spectrotemporal receptive field based models explain nearly 90% of the variance in OOR

The beat-related processing observed in the rat auditory cortex may be due to beat-specific processes, or, as we hypothesized might be more likely, due to the spectrotemporal tuning properties of recorded units. If this were the case, neural responses predicted using a standard linear-nonlinear (LN) model fitted to each unit should largely reproduce observed OORs. To test this, we first estimated each unit’s spectrotemporal receptive field (STRF), or the linear model that describes the frequency and timing properties of incoming sounds that would either excite or inhibit a neuron. Next, we estimated the unit’s static sigmoid output nonlinearity to arrive at a fitted LN model for each unit (see *Methods*). The LN model was fitted 20 times for each unit, each time using that unit’s responses to 19 of the musical excerpts while setting aside one excerpt as a test song. This ensured that predicted neural responses for a test song were true predictions since the model was not trained on the test excerpt. In this manner, firing rate predictions were generated for each unit and each musical excerpt, and these were then analyzed to arrive at predicted OOR values.

An STRF from an example unit is shown in Fig 5A, with frequency on the y-axis and stimulus history on the x-axis. This unit shows a preference for frequencies at and above 16 kHz, and is excited if sounds in that frequency range were heard 25 ms ago but inhibited if they occurred 40 ms ago. A short excerpt from a test song is shown in Fig 5B, where it can be seen that LN model predictions are in good agreement with observed firing rates. Fig 5C shows consensus OOR values for each musical excerpt based either on observed (x-axis) or predicted (y-axis) firing rates. The LN model slightly underestimates OORs (p<0.001, Wilcoxon paired signed-rank test, N = 20 songs), suggesting that there is some nonlinear process that slightly increases OOR beyond processes captured by a standard LN model. However, despite this minor difference, the LN model successfully accounts for 89% of the variance in OOR values for the tested musical excerpts (p<10^−6^, Pearson correlation, N = 20 songs). Predictions made using the linear STRF alone (without the static nonlinearity) accounted for 61% of the variance in OOR (p<0.01, Pearson correlation, N = 20 songs).

**Fig 5.**
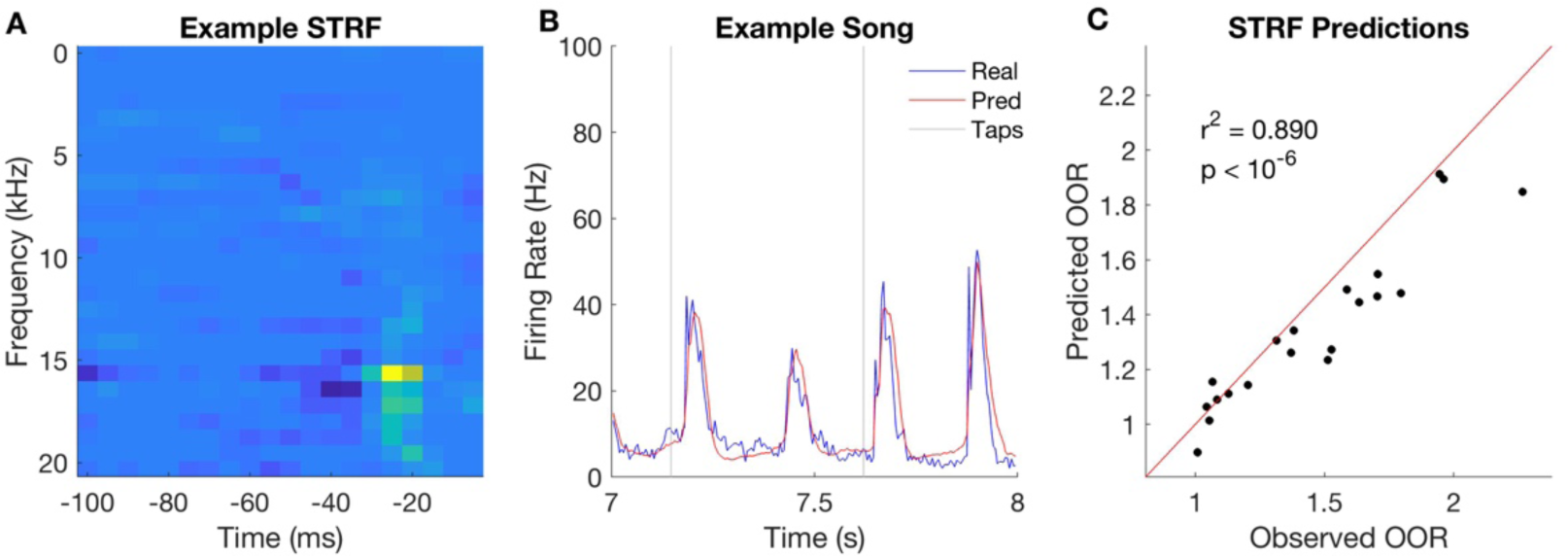
Cortical firing rate predictions based on fitted linear-nonlinear (LN) models incorporating spectrotemporal receptive fields (STRFs). **(A)** STRF from an example unit, with frequency on the y-axis and time on the x-axis and color representing the coefficients. This unit shows a classic pattern of excitation and inhibition in a relatively narrow frequency range. Convolving this filter with the spectrogram of a sound stimulus, and then applying a static nonlinearity, would result in the LN model’s prediction of this unit’s firing rate over time. **(B)** Measured (blue) and LN model predictions (red) of the population firing rate for a 1 s segment of an example musical excerpt. Gray vertical lines mark consensus tap times in this segment. **(C)** Observed (x-axis) versus predicted (y-axis) consensus on:off-beat ratios for each song. LN models account for 89% of the variance in OOR.

## Discussion

The aim of this study was to explore how firing rate transients in the auditory cortical representation of music might set the stage for the perception of musical beat. Our results, based on the twenty musical excerpts that were diverse in tempo and genre, revealed that population firing rates were on average higher on the beat than off the beat, and that large on:off-beat ratios (OORs) were a distinguishing feature of the beat interpretations most commonly tapped by human listeners. While small differences between on-beat and off-beat responses were already present in auditory nerve model responses, these differences were substantially amplified in auditory cortical responses. Furthermore, musical excerpts that evoked larger OORs in the auditory cortex also showed stronger tapping consensus among listeners. Finally, the spectrotemporal receptive field (STRF) properties of cortical units were able to account for the magnitude of the OOR each musical excerpt would induce. Together, these findings suggest that large OORs in the auditory cortex, which arise due to the spectrotemporal tuning properties of neurons, may be key to establishing the location and clarity of the perceived beat.

It is worth noting is the extent to which the physiology corresponded to tapping behavior and the extent to which standard LN STRF models could capture the physiology for real musical excerpts. These observations strongly suggest that the related low-level mechanisms of neuronal adaptation^10^, amplitude modulation tuning^30^, and STRFs play a formative role musical in beat perception. This is not inconsistent with the theory that the induction of the beat percept is the result of an interaction between “bottom-up” sensory processes and “top-down” cognitive ones^31^. Our data suggest that beat perception may really begin weakly at the ear, with neural activity showing stronger correspondences to behavior as information ascends through the brainstem and primary cortical structures of the ascending auditory pathway^32,33^. Since these parts of the ascending auditory system are often highly conserved across mammalian species^34–37^, cross-species investigations may be a promising way to understand the neural signals and dynamics that underlie beat induction, which to date remain mysterious.

Though our results indicate that beat perception is strongly influenced by basic physiological mechanisms and therefore only partly culturally determined, they do not imply that “bottom-up” processes could possibly explain everything. For example, some well-studied constraints on-beat perception include the tendency to perceive a beat within a frequency range of roughly 0.5–4 Hz^38^ with a special preference for 2 Hz^39^, and an overall preference for binary (e.g. 2, 4) meters over ternary (e.g. 3, 6) or other complex meters^38,40^. These constraints are likely driven by top-down influences or may result from auditory-motor interactions^8,17–25^ and are unlikely to be explained by bottom-up sensory processing alone. Furthermore, the perceived beat and its neural signatures can be modulated at will by top-down attention or mental imagery of beat structure^12,41–43^. Bringing these ideas together, we propose that the perception of beat relies on the application of learned and implicit rhythmic priors^6,7^ onto an ascending sensory representation^10,30^ with a bias towards configurations that maximize the difference between neural activity on and off the beat.

That we see as much correspondence as we do between the representation in auditory cortex and beat perception could be an indication that neural activity in the auditory cortex is a key interface between the sensory and motor and/or cognitive processes involved in beat perception. Probing the cortico-basal ganglia-thalamo-cortical loop^44^ may be a promising avenue for future investigations. Projections from auditory cortical fields to the basal ganglia have been well-characterized^45^, and the basal ganglia in humans have been repeatedly implicated in beat perception^22,43,46,47^ as well as other auditory cognitive abilities^48^. We speculate that large firing rate transients in the auditory cortex, observed in this study to co-occur with the perceived beat, could set into motion the dynamics of this loop and thereby enable the possible entrainment of cortical oscillations to the beat^21,42,49–51^. We suggest caution, however, as there is currently some debate around what constitutes neural entrainment to auditory rhythms^52–54^, and whether frequency-domain representations of rhythms and brain signals necessarily reflect beat perception^55^.

The extent of the correspondence observed in this study between auditory cortical activity in rats and human beat perception also invites the intriguing question of whether rodents too can perceive musical beat. Preliminary evidence suggests that rats can be trained to discriminate isochronous rhythms from non-isochronous ones^56^. Mice too appear capable of performing a synchronization-continuation task, and in that study, primary auditory cortex was implicated as being necessary for the generation of anticipatory motor actions^57^. These studies at minimum suggest that rodents have the capacity to perceive temporal structure and execute motor actions timed to an external isochronous rhythm. Future behavioral studies are needed to explore the limits of sensorimotor synchronization in rodents.

At the other end of the spectrum are humans, whose ability to synchronize with an external rhythm, whether it is to a metronome or to the beat in music, is spontaneous^1^, highly anticipatory^58^, innate^59^, and often involuntary^2,60,61^. The gradual audiomotor evolution hypothesis posits that the ability to entrain movements to musical beat relies on strong coupling between the auditory and motor systems, and that the neurophysiology and behavioral capacity to do so evolved gradually^20^. This hypothesis is supported by evidence that nonhuman primates, like humans, are capable of producing tempo-flexible anticipatory movements in time with a metronome^62,63^ and can detect rhythmic groupings, but cannot detect or synchronize to a musical beat^4^. The dissociation between perceiving auditory rhythms and perceiving musical beat may relate to findings that distinct networks underpin “duration-based” and “beat-based” temporal predictions^64–66^. It is important for future studies in the area of beat perception to be clear about precisely what is being perceived, since there is demonstrable nonequivalence between the detection of a pulse in isochronous rhythms, a pulse in real music, and beat in the context of the different levels of nested hierarchical structure present in music, the latter of which has arguably not yet been demonstrated in any nonhuman species^67^.

This leads to the question of why beat perception exists in the first place. Some clues might be found in parallels that beat perception has with other abilities, particularly with the human capacity for language^68–70^. Another possibility is that beat may provide a way to quickly assess locomotion speed from the sound of a complex gait. Though this speculation has not yet been tested directly, gait studies have shown that humans are able to assess a number of attributes of a walker based only on their walking sounds, including gender, posture, and emotional state^71,72^.

However, at the heart of these complex abilities are neural circuits that are very old and also underlie more general auditory cognitive abilities^73^ such as perception of time^74^ and prediction of future sensory inputs^75^. Therefore, a unified perspective that would bring all of this together is that the information processing performed by the auditory system up to primary auditory cortex is largely consistent across most mammals, but the complexity of the operations the organism ecologically needs to perform with this information may be the determinant for what is “top-down.” Our data suggest that strong firing rate transients in the neural representation of real music may shape where the beat is felt, and while an on-beat neural emphasis is certainly not the whole story, it is a lead worth exploring further. Ultimately, this work underscores the importance of low-level auditory processing in creating a representation of sound where certain features are emphasized based on temporal context, a representation on which other high-level processes rely to give rise to complex perception.

## Methods

### Stimuli

The 20 songs tested were the training dataset for the MIREX 2006 beat tracking algorithm competition^76^. Each song had beat annotations collected from 40 human listeners^28^. Only the first 10 s of songs and beat annotations were used in this study.

### Surgical Protocol

All procedures were approved and licensed by the UK home office in accordance with governing legislation (ASPA 1986). Three female Lister Hooded rats weighing approximately 250 grams were anesthetized with an intraperitoneal injection of 0.05 ml domitor and 0.1 ml ketamine. To maintain anesthesia, a saline solution containing 16 ug/kg/h domitor, 4 mg/kg/h ketamine, and 0.5 mg/kg/h torbugesic were infused continuously during recording at a rate of 1 ml/h. A craniotomy was performed 4.7 mm caudal to bregma and extending 3.5 mm lateral from the midline on the right hand side.

Recordings were made using a 64 channel silicon probe (Neuronexus Technologies, Ann Arbor, MI, USA) with 175 um^2^ recording sites arranged in a square grid pattern at 0.2 mm intervals along eight shanks with eight channels per shank. The probe was inserted into the auditory cortex in a medio-lateral orientation wherever possible.

The 20 songs were played in randomized order for a total of 12 repeats, with 3 seconds of silence separating each song from the next. Stimuli were presented binaurally through headphones at 80 dB SPL. Sounds were presented with a sampling rate of 48828.125 Hz, and data were acquired at a sampling rate of 24414.0625 Hz using a TDT system 3 recording setup (Tucker Davis Technologies).

## Data Analysis

### Tapping Analysis

To calculate consensus tap times, the histogram of tap times, pooled across the 40 subjects and then binned using 2 ms bins, was smoothed using a Gaussian kernel with a width (standard deviation) of 40 ms. This width was chosen because visual inspection of tap histograms showed the standard deviation around taps to be approximately 40 ms, so a Gaussian kernel with that width would approximate a “matched filter.” The precise width of the smoothing kernel was not critical to our results as long as it roughly matched the spread in the data. A peak-finder (*findpeaks.m*, built-in Matlab function) was then used to identify peaks that were larger than 40% of the maximum value in the smoothed histogram. The consensus inter-tap-interval (ITI) for a song was taken to be the mean interval between successive peaks, after the exclusion of intervals larger than 1.5 times the median inter-peak-interval (which would happen if the peak-finder missed a peak). The consensus phase was determined by finding the offset that optimally aligned a temporal grid with consensus ITI spacing with the peaks found by the peak-finder. Consensus tap times can be described by a consensus ITI (beat period) and consensus offset (beat phase) combination for each song.

On-beat neural activity was defined as the average population firing rate in the 100 ms following consensus tap times, and off-beat neural activity was the average population firing rate during all time excluding these on-beat windows. The justification for this definition is that (i) the true perceived beat location is almost certainly just after a listener taps, given the well documented tendency of listeners to anticipate the beat with their movements by several tens of milliseconds (negative beat asynchrony)^61^, (ii) defining off-beat activity as all neural activity that is not on the beat is consistent with previous work^10^, and (iii) an interval of 100 ms is less than one half a beat cycle for the fastest beat period observed in these data of 273 ms. The precise choice of time window is not critical, and this was confirmed by running all analyses using on-beat windows that ranged between 40 ms and 120 ms in 10 ms increments. The results were entirely consistent with those presented here for a time window of 100 ms, and if anything, slightly stronger when shorter time windows were used.

To compute the strength of the consensus, an “ideal tap histogram” was constructed by assuming all 40 listeners tapped precisely at each consensus beat time as determined by the excerpt’s consensus ITI and phase. A realistic degree of motor error was added by convolving this with a Gaussian kernel whose width was 5% of the beat period. The same 5% Gaussian kernel was then used for kernel density estimation on the two signals: the raw pooled histogram of (measured) tap times that already contained motor error, and the idealized tap histogram with motor error added. The choice of temporal filter value was guided by the magnitude of errors reported in studies of human sensorimotor synchronization^77–79^, but other kernel widths close to 5% also produce consistent results. The correlation coefficient between real and idealized tap density estimates for a given musical excerpt was taken as a measure of the strength of the tapping consensus, where a large value would indicate a high degree of similarity between real and “ideal” tapping behavior. Estimation of the real and idealized tap densities is also possible using a constant width (e.g. 40 ms) Gaussian kernel rather than a proportional one. However, while doing so would lead to the to the same main result shown in Fig 4, this measure of tapping consensus strength would have the undesirably effect of also correlating with song tempo since, as mentioned above, it is well-established that the magnitude of sensorimotor synchronization errors scale with interval duration.

### Electrophysiology Data Preprocessing

Offline spike sorting and clustering was done on the raw data using an automated expectation-maximization algorithm (Spikedetekt/Klustakwik)^80^, and clusters were manually sorted using Klustaviewa (Cortical Processing Lab, University College London). Firing rates over time for multi-units were calculated by binning spike times into 5 ms bins, which resulted in peri-stimulus time histograms (PSTHs) at an effective sampling rate of 200 Hz.

To determine whether spikes were reliably stimulus-driven, a noise power to signal power cutoff of 40 was chosen^81^. Song 1 was arbitrarily chosen to the be the stimulus for which the repeatability of responses was measured. Units that failed to show a noise power to signal power ratio less than 40 based on the 12 repeats were excluded from further analysis, leaving a total of 98 multi-units. All subsequent analyses were performed using custom-written Matlab code.

### Fitting the LN Model

The relevant scripts used at all stages of this process are available on Github^82^. First, music stimuli were transformed into a simple approximation of the activity pattern received by the auditory pathway by calculating the log-scaled spectrogram (‘cochleagram’)^82–84^. For each sound, the power spectrogram was taken using 10 ms Hanning windows, overlapping by 5 ms. The power across neighboring Fourier frequency components was then aggregated using overlapping triangular windows comprising 27 frequency channels with center frequencies ranging from 50 Hz to 20,319 Hz (1/3 octave spacing). Next, the log was taken of the power in each time-frequency bin, and finally any values below a low threshold were set to that threshold. These calculations were performed using code adapted from melbank.m (http://www.ee.ic.ac.uk/hp/staff/dmb/voicebox/voicebox.html). The STRF model was trained to predict the firing rate at time *t* from a snippet of the cochleagram extending 100 ms (20 time bins) back in time from time *t*. The linear weights describing the firing rate of each neuron were estimated by regressing, with elastic net regularization, each neuron’s firing rate at each time point against the 100 ms cochleagram snippet directly preceding it. Regularization strength was set by using a randomly chosen 10% of time bins from the cross-validation set as a validation set, and then by choosing the regularization parameters that led to the fit on the validation set with the lowest mean squared error. A sigmoidal nonlinearity^85^ was then fitted to map from the linear activation to the predicted PSTH such that it minimized the error between the predicted PSTH and the observed PSTH. LN model predictions of a unit’s PSTH to a test song were made by first convolving the cochleagram of the test song with the linear STRF and then applying the nonlinearity. Each unit’s LN model was calculated 20 times, each time setting a different song aside as the test set. This was done so that PSTH predictions for any musical excerpt were true predictions since that excerpt was not included in the training set for the model.

## Supporting information

