## Supplementary material for "Beat-Relevant Signals in Auditory Cortical Responses to Musical Excerpts"

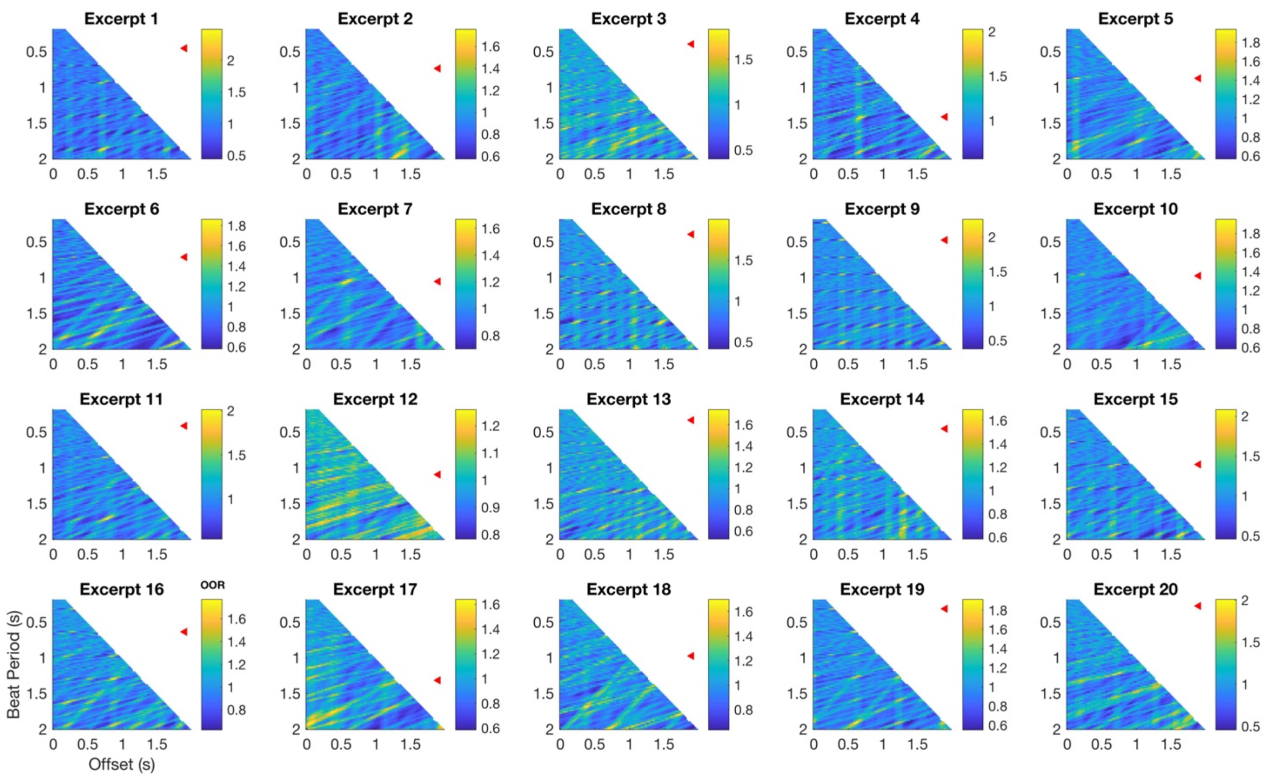


**Fig S1. Heatmaps illustrating hypothetical OORs for a range of possible beat structures based on auditory cortical firing rates.** Each panel is one excerpt, as labelled in the MIREX 2006 database. Colors show OOR values for beat period (y-axis) and beat phase (x-axis) combinations between 200 ms and 2 s (or tap rates of 5 Hz down to 0.5 Hz) in 0.02 s increments. Red triangles mark the row corresponding to that song's consensus beat period. See Fig 2A in the main text.

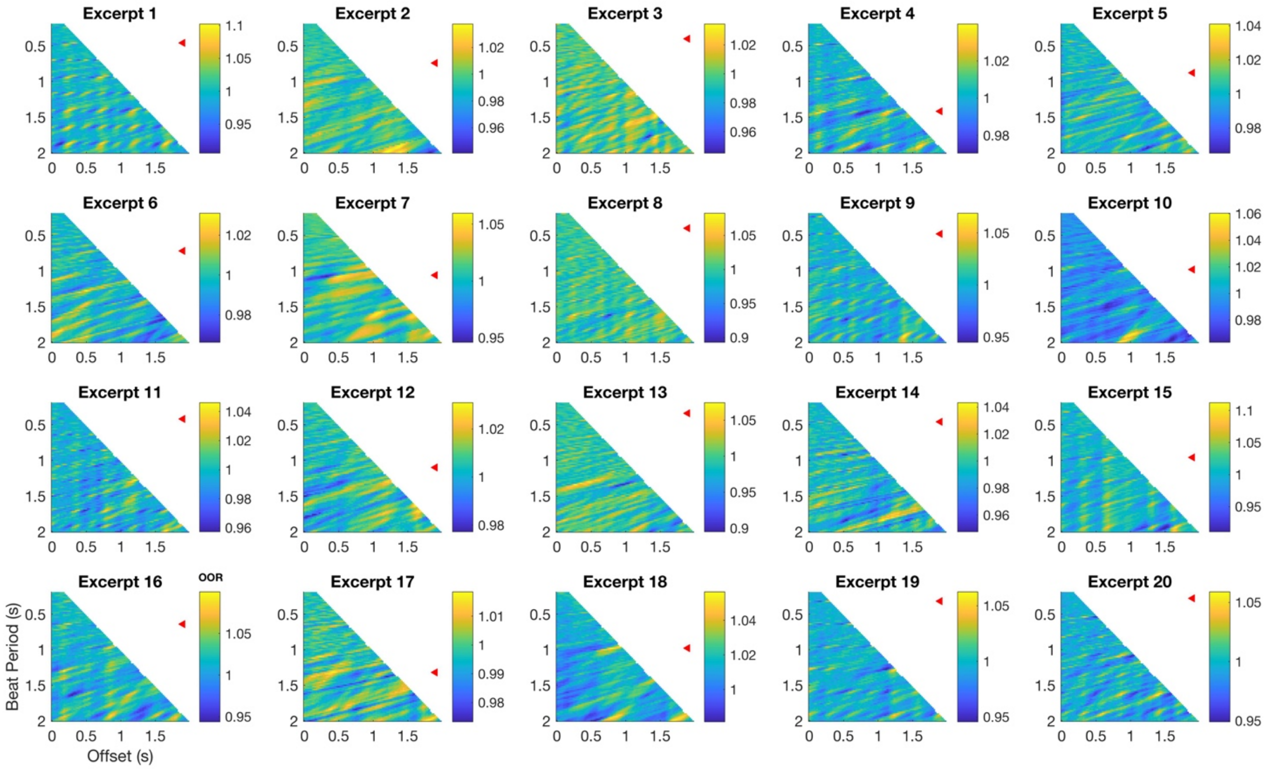


**Fig S2. Heatmaps illustrating hypothetical OORs for a range of possible beat structures based on auditory nerve model.** Each panel is one musical excerpt, as labelled in the MIREX 2006 database. Colors show OOR values for beat period (y-axis) and beat phase (x-axis) combinations between 200 ms and 2 s (or tap rates of 5 Hz down to 0.5 Hz) in 0.02 s increments. Red triangles mark the row corresponding to that song's consensus beat period. Note the difference in range of OORs compared to cortical data in Fig S1. See Fig 2B in the main text.

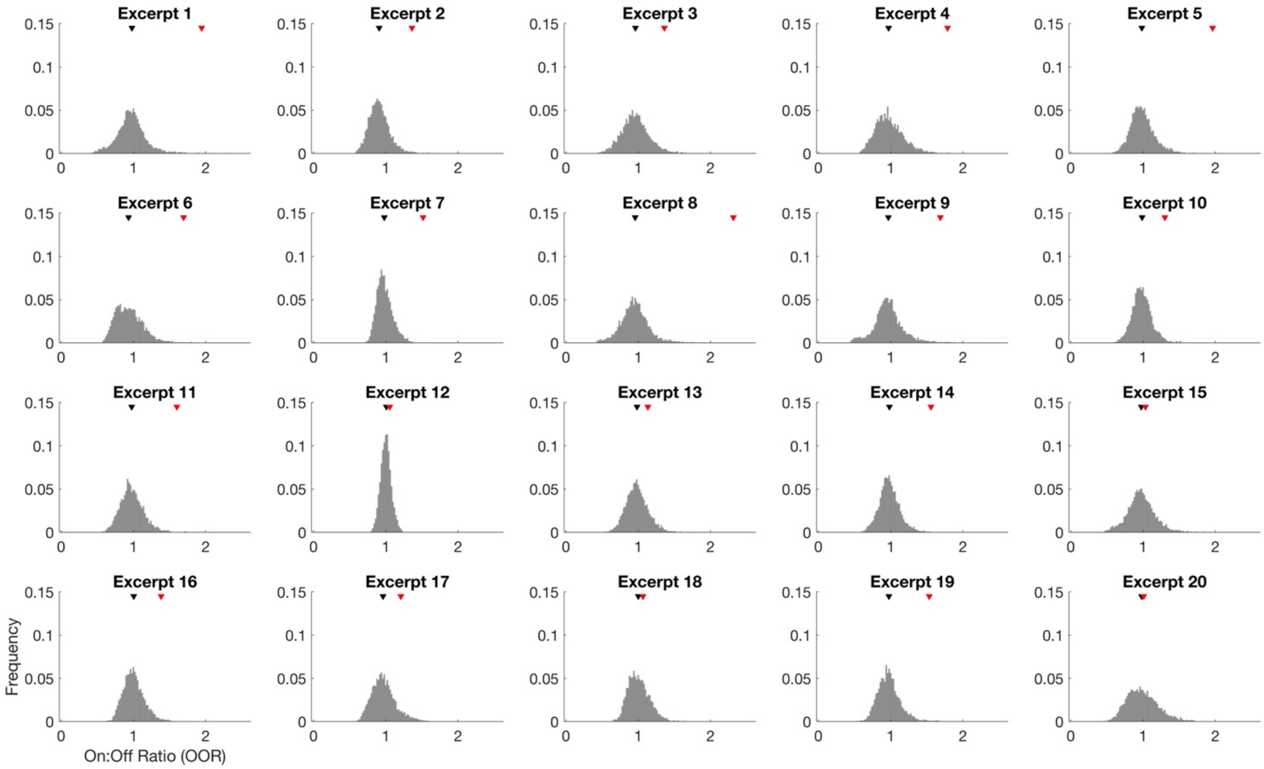


**Fig S3. Distribution of hypothetical OOR values in the auditory cortex for each song.** Each panel is one musical excerpt, as labelled in the MIREX 2006 database. Histograms show OOR values at all sampled hypothetical beat period and beat phase combinations for a given piece of music. The gray triangle shows the median of this distribution, and the red triangle shows the song's consensus OOR value. The consensus OOR is high relative to the underlying distribution of OORs for most songs. See Fig 2C in the main text.

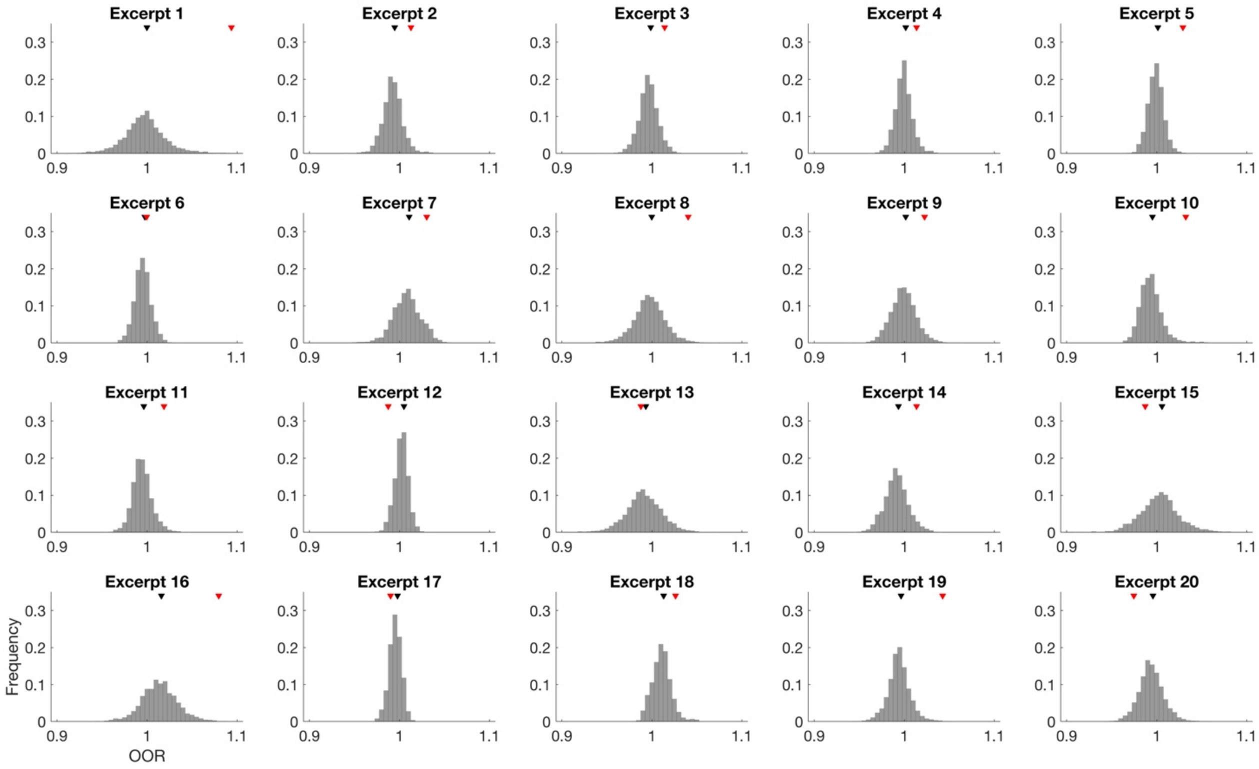


**Fig S4. Histograms of max OOR values based on auditory nerve model.** Each panel is one musical excerpt, as labelled in the MIREX 2006 database. Histograms show OOR values at all sampled hypothetical beat period and beat phase combinations for a given piece of music. The gray triangle shows the median of this distribution, and the red triangle shows the song's consensus OOR value. Note the substantially narrower range of OORs (x-axis) compared to Fig S3. See Fig 2D in the main text.

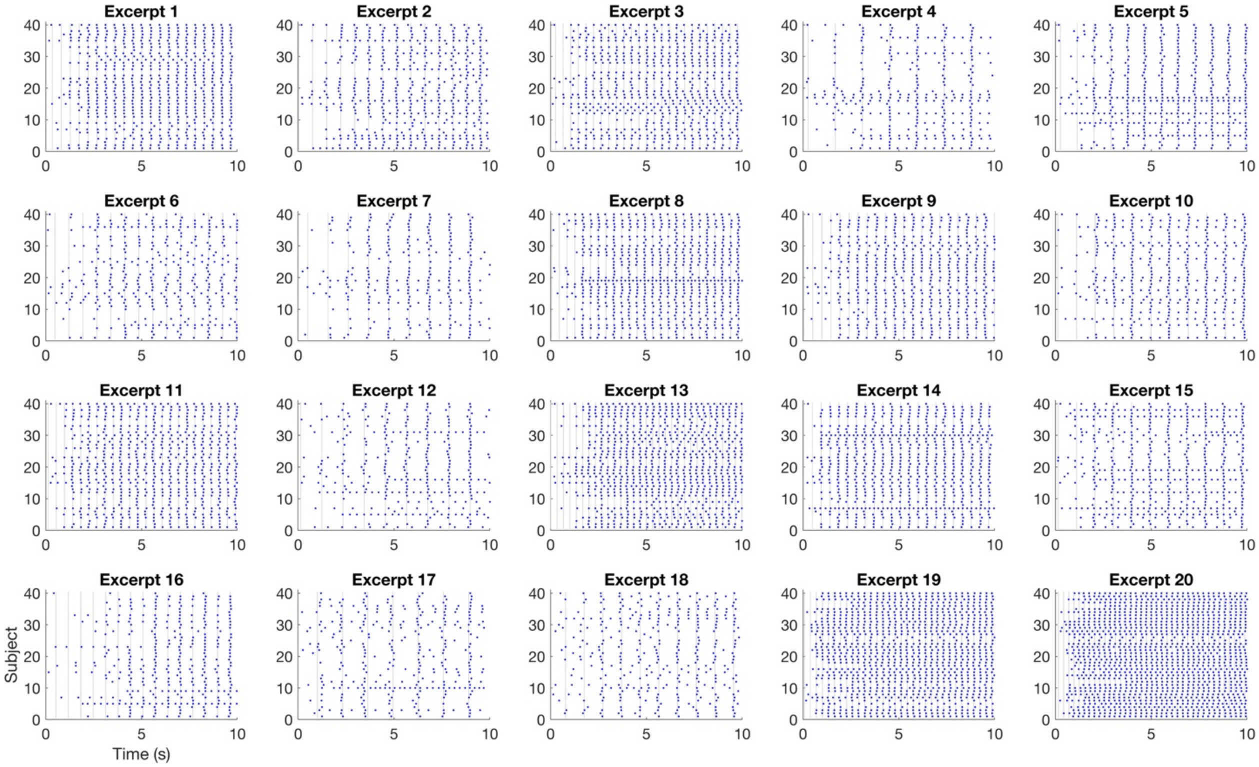


**Fig S5. Tap rasters for all songs.** Each panel is one musical excerpt, as labelled in the MIREX 2006 database. Each dot is one tap, each row is one subject and the position of taps along the x-axis represents when that subject tapped. See Fig 3 in the main text.

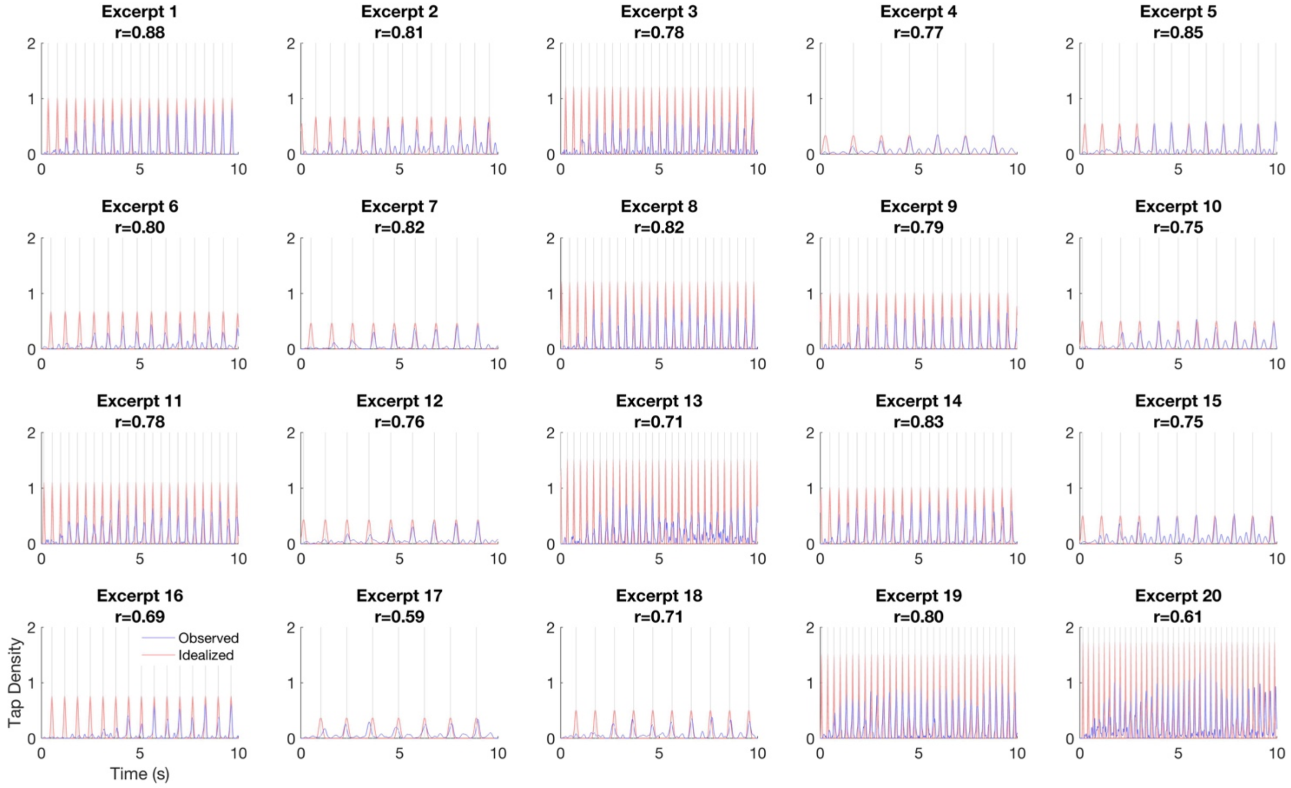


**Fig S6. Observed and idealized tap density estimates.** Each panel is one musical excerpt, as labelled in the MIREX 2006 database. Tap density estimates are based on tap times pooled across subjects, binned with 2 ms bins, and smoothed with a Gaussian kernel with a standard deviation of 5% of the consensus beat period (blue). Shown in red is a tap density estimate of the "ideal" tap histogram (with realistic motor error) that would have been obtained if all subjects had tapped on every consensus beat (see *Methods*). The correlation coefficient (r) between real and idealized tap density was taken as a measure of each excerpt's tapping consensus. See Fig 3 in the main text.
